# Towards a Clinically-based Common Coordinate Framework for the Human Gut Cell Atlas - The Gut Models

**DOI:** 10.1101/2022.12.08.519665

**Authors:** Albert Burger, Richard Baldock, David J Adams, Shahida Din, Irene Papatheodorou, Michael Glinka, Bill Hill, Derek Houghton, Mehran Sharghi, Michael Wicks, Mark J Arends

## Abstract

**Background:** The Human Cell Atlas resource will deliver single cell transcriptome data spatially organised in terms of gross anatomy, tissue location and with images of cellular histology. This will enable the application of bioinformatics analysis, machine learning and data mining revealing an atlas of cell types, sub-types, varying states and ultimately cellular changes related to disease conditions. To further develop the understanding of specific pathological and histopathological phenotypes with their spatial relationships and dependencies, a more sophisticated spatial descriptive framework is required to enable integration and analysis in spatial terms.

**Methods:** We describe a conceptual coordinate model for the Gut Cell Atlas (small and large intestines). Here, we focus on a Gut Linear Model (1-dimensional representation based on the centreline of the gut) that represents the location semantics as typically used by clinicians and pathologists when describing location in the gut. This knowledge representation is based on a set of standardised gut anatomy ontology terms describing regions *in situ*, such as ileum or transverse colon, and landmarks, such as ileo-caecal valve or hepatic flexure, together with relative or absolute distance measures. We show how locations in the 1D model can be mapped to and from points and regions in both a 2D model and 3D models, such as a patient’s CT scan where the gut has been segmented.

**Results:** The outputs of this work include 1D, 2D and 3D models of the human gut, delivered through publicly accessible Json and image files. We also illustrate the mappings between models using a demonstrator tool that allows the user to explore the anatomical space of the gut. All data and software is fully open-source and available online.

**Conclusions:** Small and large intestines have a natural “gut coordinate” system best represented as a 1D centreline through the gut tube, reflecting functional differences. Such a 1D centreline model with landmarks, visualised using viewer software allows interoperable translation to both a 2D anatomogram model and multiple 3D models of the intestines. This permits users to accurately locate samples for data comparison.

## Background

Following a short preamble introducing the Human Cell Atlas endeavour, the main objective of this background section is to provide the reader with the biomedical context of our work. Specifically, we begin with brief introductions of 1) human gut anatomy, the focal point of our models, 2) Inflammatory Bowel Disease, a primary medical concern ultimately to benefit from this research, 3) clinical investigations, which underpin the nature of our models, and 4) single Cell RNA sequencing technology, which is the main development pushing biomedical atlasing work down to the cellular level. Building on this basis, we set out the general case for so-called Common Coordinate Frameworks and the specific objectives of our work.

### Introduction to Human Cell Atlas

The mission of the Human Cell Atlas (HCA) programme [1] is “To create comprehensive reference maps of all human cells—the fundamental units of life—as a basis for both understanding human health and diagnosing, monitoring, and treating disease.” [2] The human body is a complex amalgamation of cells organised into tissue, organs and systems which can be studied in health and disease states. The ability to study complex organisms at the most basic cellular level has generated vast quantities of molecular data necessitating suitable data capture and modelling platforms to support the interpretation of the data. The ability to visualise and map cell to tissue and tissue to organ data will allow in future for a more comprehensive understanding of changes related to health and pathological conditions.

### Human Gut Anatomy

The gastrointestinal tract can be represented as a long cylindrical tube from oesophagus through stomach, small intestines, large intestines, to anal canal, terminating at the anus. The main function of the gut is to digest and absorb nutrients with the excretion of waste products. It also has essential roles in endocrine, immune and barrier function, delicately balancing the symbiotic relationship with the microbiome and supporting continuous epithelial tissue renewal. Here, we focus on the small and large intestines, from gastro-duodenal junction to anus. These gut components have internationally standardised gut anatomy ontology terms that describe the various regions, such as duodenum, jejunum and ileum of the small intestines, and caecum, ascending colon, transverse colon, descending colon, sigmoid colon, rectum and anal canal of the large intestines. A number, but not all, of the junctions between these component regions are separated by established landmarks, such as ileo-caecal valve, hepatic flexure, splenic flexure and anorectal junction for example. These gut regions with landmarks, together with consensus average length measurements, can be used to generate a 1-dimensional map or model of the gut that allows normal or disease samples to be located more precisely.

### Inflammatory Bowel Disease

Mapping of disease location accurately within the gut is important for Inflammatory Bowel Diseases (IBD). These are chronic inflammatory conditions of the gastrointestinal tract with an increasing incidence worldwide [3]. The underlying inflammation is postulated to be secondary to the interactions between the microbiome, an activated immune system and mucosal barrier dysfunction in genetically susceptible individuals. The increase in incidence has been linked to adoption of a westernised diet with ultra-processed food [4, 5] and medications such as proton pump inhibitors [6, 7]. There are two main types of IBD: Ulcerative Colitis and Crohn’s Disease. Ulcerative Colitis affects the large bowel, often starting in the rectum and progressing proximally, resulting in abdominal pain and a change in bowel function. Crohn’s Disease is the more complex disorder affecting any part of the gastrointestinal tract from mouth to anus, with distinct disease manifestations associated with the specific region of the affected gut [8].

### Clinical Investigations

IBD is diagnosed by standard methods including clinical assessment, radiological, endoscopic and histological evaluation. The Lennard-Jones criteria are considered the gold standard for confirming the diagnosis of Crohn’s Disease [9]. Endoscopic evaluation is used frequently to obtain tissue samples which are analysed to confirm the pathognomonic changes of discontinuous transmural inflammation, often with a fissuring pattern of deep ulceration and fibrosis [10]. In addition to the fissuring ulcers, there is both acute and chronic inflammation with focal cryptitis, crypt destruction and granuloma formation in around 60% CD cases. Longstanding inflammation may predispose to dysplasia which in some may evolve to invasive adenocarcinoma. Some patients may develop fibrotic tissue resulting in narrowing or stricturing of the affected bowel precipitating bowel obstruction. Subsequently, fibrotic tissue can be surgically excised although the mechanism of fibrosis is poorly understood and other treatment strategies are less effective [11].

It is a challenge to accurately map these changes to their correct positions in a three-dimensional model to illustrate the distribution within the gastrointestinal tract. If the patient has undergone surgery and there is associated radiological imaging, then location can be determined reasonably straightforwardly. During endoscopic procedures (and many surgical resections), the endoscopist (or surgeon or pathologist) usually describes the small or large intestinal region (e.g. ileum, ascending colon, etc) involved by a lesion and sometimes provides a distance (in cm) from the anus using the distance markings on the endoscope surface (see Figure 1), or for surgical resection specimens, a distance of the lesion from a landmark such as the ileo-caecal valve or the resection margin of the specimen. Some change to clinical practice is required to capture these data *routinely* as distances from recognisable gut landmarks.

**Figure 1:**
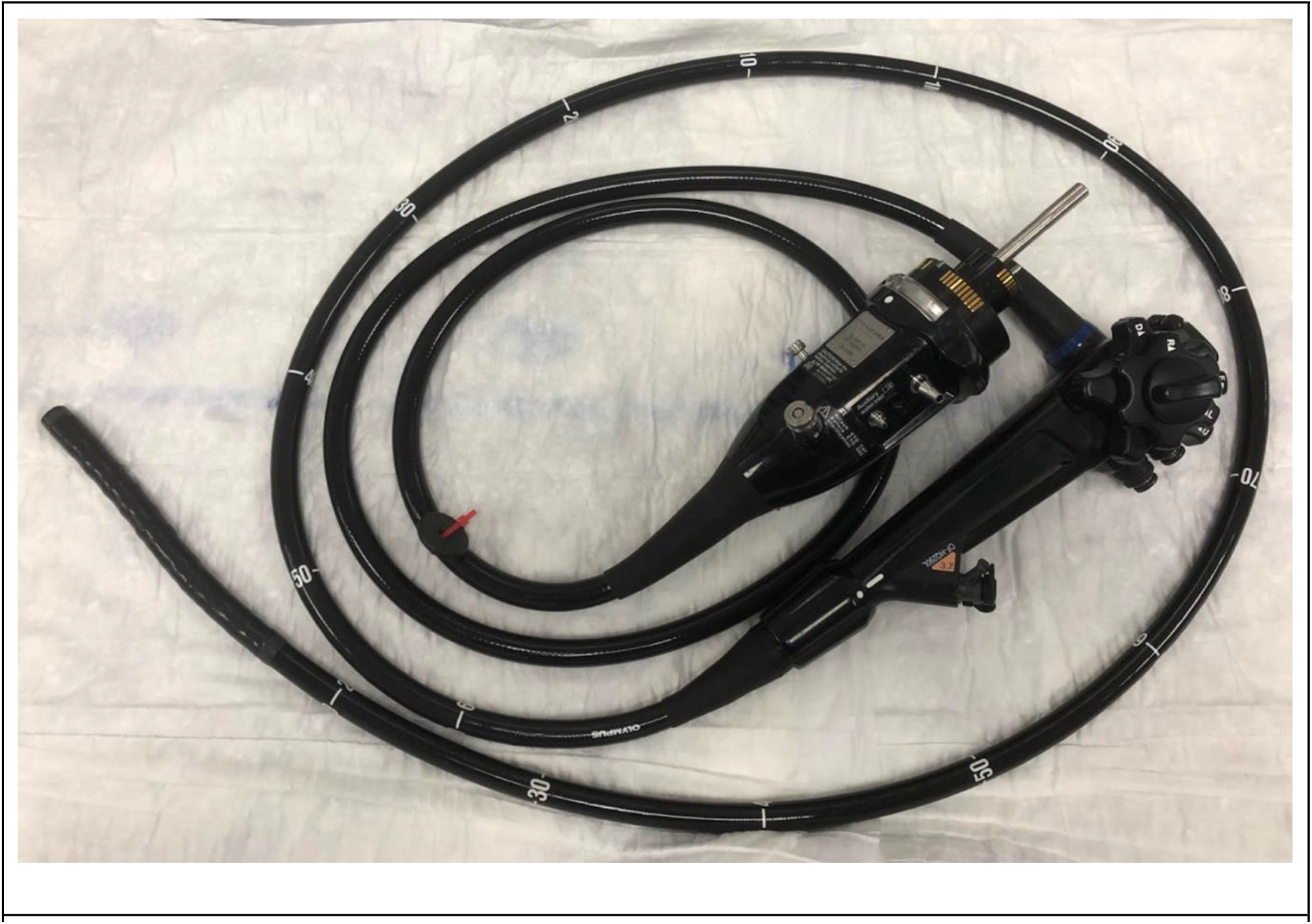
Endoscope showing distance markings. These could be used to establish a gut-location by taking readings for the sample location coupled with the nearest proximal and nearest distal gut landmarks e.g. the hepatic and splenic flexures for locations within the transverse colon.

### Single Cell RNA Sequence Data Analysis

Single-cell sequencing is a powerful technology for profiling the transcriptome of large numbers of individual cells (see [12], [13] and [14] for a recent review). The technique generates large amounts of data that requires specialised computational and statistical analysis. Generally, single cells are isolated into wells of a plate or into droplets, such that transcripts from each cell can be barcoded or tagged (marking them with a unique molecular identifier; UMI), allowing the expression profile of the cell to be ascertained after RNA-sequencing, which is normally performed on pools of transcriptomes from many cells. The major variables in all single cell sequencing experiments are the number of cells profiled and the depth of sequence generated for each cell. The initial steps of single cell sequencing data quality control include removing data associated with UMIs that are not well represented, these are often associated with cells that are dying or are damaged, and to then examine the proportion of multi-mapping, un-mappable and mitochondrial reads for each cell, the frequency of which tend to correlate with poor data quality. Since the aim is to profile the transcriptome at single cell resolution, empty droplets, cell-free RNA and doublet-cells are removed using software such as EmptyDrops, SoupX and DoubletFinder respectively. Following on from these steps the data is normalised to account for differences in sequencing depth and where appropriate batch correction to account for non-biological factors such as time of sample collection. Further data processing steps can involve data smoothing and imputation, cell cycle analysis, unsupervised clustering as a prelude to dimensionality reduction and data visualisation, which can be performed using approaches such as PCA, t-SNE and UMAP. Where differential expression analyses are a key parameter, various methods have been developed including MAST and MetaCell. The field of single cell sequence analysis is rapidly evolving with many robust and elegant approaches allowing data exploration and there is a requirement for integration of such data with histological, radiological, clinical disease metadata and other data using a common coordinate framework approach. In addition, mapping the locations of the source tissue samples within the gut context will allow the discovery and analysis of the gradients of variation along the proximal-distal gut axis and reveal novel understanding of the gut biology. Without a mechanism for capturing gut location this aspect of gut-biology will remain undiscovered.

### Towards a Human Gut Cell Atlas Common Coordinate Framework

The primary aim of the Human Gut Cell Atlas (HGCA) is to capture a detailed atlas of tissue and single cell data in the spatial context of the adult human gut. Whether for clinical purposes for individual patients or more general research studies concerning the gut, data ranging from patient-specific information, histopathological image data and radiological images, to single cell sequencing data of gut cells as part of research work, all are now collected and stored by hospitals and research institutions, respectively. Data integration is a key prerequisite to facilitate AI techniques, in particular Machine Learning, to derive medically useful knowledge from these large, distributed data sources, in order to reveal the spatial organisation of the underlying molecular and cellular processes in normal and diseased samples. One of the primary integration criteria is the anatomical origin of tissues and cells which the collected data refers to. In the context of the Human Cell Atlas, such anatomical locations are to be recorded using a computational framework called the *Common Coordinate Framework (CCF)* [15].

The number and types of use cases for a Human Gut Cell Atlas and therefore the requirements for its CCF, are large and varied. A balance has to be struck between catering for all eventualities and a simplicity that makes the use of the CCF practical. In this paper, we describe a CCF for the human gut that is based on clinical practice, has at its core an easily understandable 1D gut model, but extends to complex 2D and 3D representations. A mechanism for capturing proximal-distal gut location is critical to enable not just an atlas of cell-types as revealed by scRNA-seq but also the gradients of change of the cell-types, sub-types and cell states, as well as cell populations along the gut axis and how that links to gut anatomy in both health and disease. Once we have introduced our own models for a Human Gut Cell Atlas CCF in the Methods and Results sections, we provide further details on related frameworks in the Discussion.

### Objectives

The primary objective of the Human Gut Cell Atlas programme is to enable data integration of all data-types to deliver a research and analysis capability to support science discovery and clinical benefit in the context of the gut and related tissue diseases and pathological abnormalities. Our objective with this work is to deliver a practical CCF for the gut that make this possible. For that we have developed a conceptual model of the gut based on the natural coordinate of distance along the gut midline with semantic extension to specific tissues and cells. In addition, we develop a mapping mechanism that allows cross comparison with 2D and 3D gut representations including patient-specific data. A further aim is the interoperability of the proposed CCF with other similar efforts, to facilitate cross-CCF data integration.

In the Methods section we set the scientific context for our models in terms of the specific use case underpinning our work and then describe how the models were developed. The Results section presents the 1D, 2D and 3D models that we created and which form the core of the proposed CCF. In addition, a publicly accessible online tool illustrating the use and interaction of these models is presented. Related work is reviewed in the Discussion section, as are limitations as well as future prospects of the Gut CCF. The Conclusions summarise the primary contribution of the models and their implementations, as well as their importance and potential impact in the context of the Human Gut Cell Atlas endeavour.

## Methods

### Edinburgh-Cambridge Helmsley Trust project HGCA CCF use-case

Although the exact CCF requirements across different projects will vary, the project described here includes many of the typical components for this kind of work, and thus facilitated the development of the gut models with common clinical and research practice in mind. For this project, Crohn’s disease lesion samples are collected from surgical resections. From the *resection specimen* tissue *slices* are taken from various sample points capturing both diseased and morphologically healthy (no visible pathological abnormality) tissue. The CCF must be able to capture the location from where in the gut the samples (either biopsies or blocks from a surgical resection) were taken, and for multiple tissue slices, the relative location of slices in terms of their sequence order as taken from the surgical resection specimen. Following slicing of a surgical resection specimen of gut, one or more parts of the slices form blocks of tissues for further processing, for either dissociation of fresh tissue into single cells for single-cell transcriptome sequencing, or fixation for histological analysis. In the latter case tissue blocks are fixed in buffered formalin, processed into paraffin and *sections* are cut for staining, scanning and analysis. Both the source of the sections – in terms of their original blocks – and their relative ordering and adjacency in the blocks must be tracked. Histology and sequence data generated during analysis are annotated with relevant CCF location information to allow the integration of data based on the same precise location of tissue, but also to map across different samples from different patients. Datasets will be made available where possible in accordance within the appropriate legal framework or within a secure research environment [16]. A conceptual overview of the project is provided in Figure 2.

**Figure 2:**
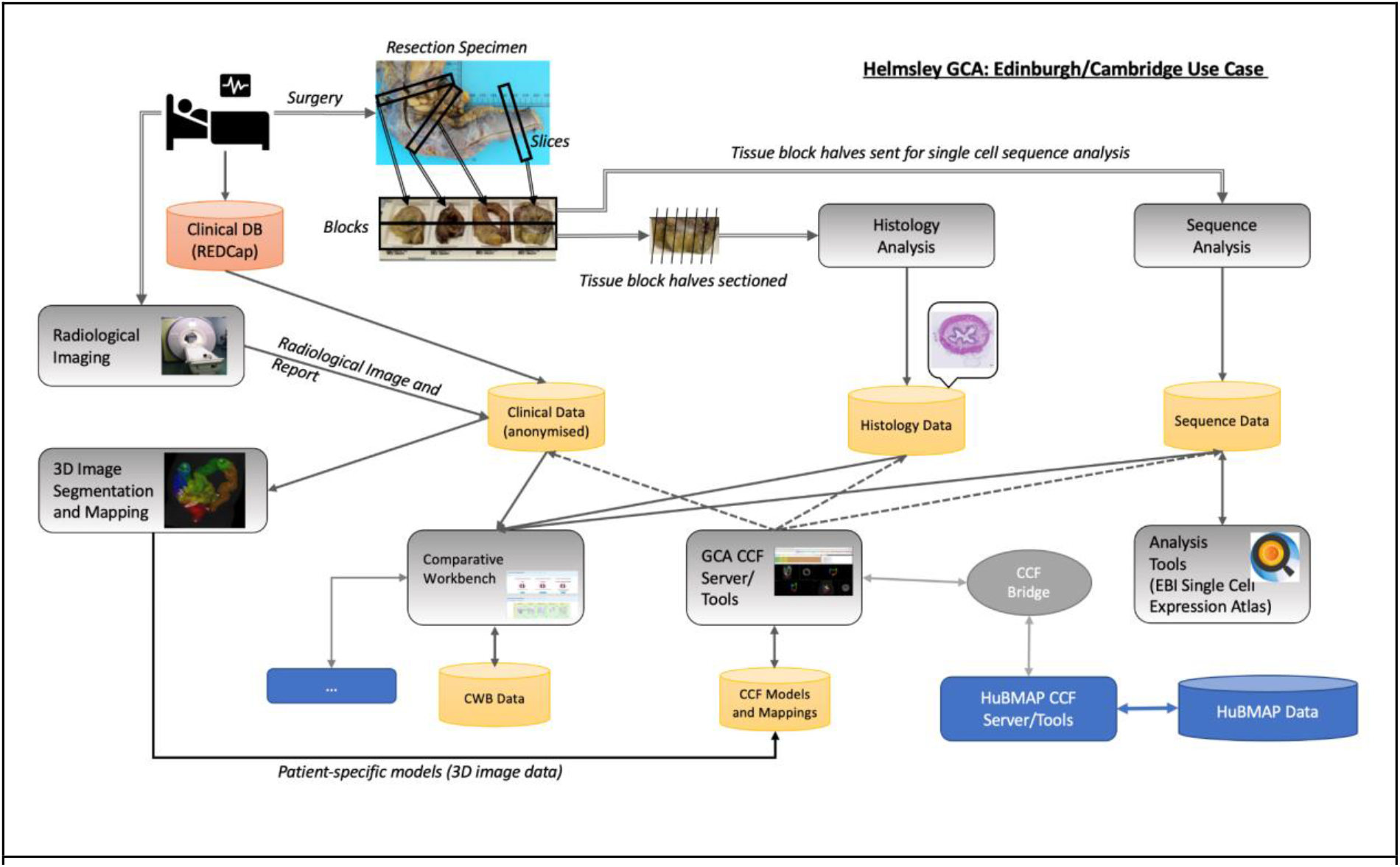
Edinburgh-Cambridge Helmsley Trust project HGCA CCF use-case. The project design includes pre-surgery MRI imaging followed by Crohn’s lesion resection. The resected material is processed to produce histological sections for staining and imaging matched to adjacent tissues used for single-cell transcriptomics. The transcriptome data is archived at and accessible from the Single Cell Expression Atlas and the radiological and histology image data is submitted to the HCA archive.

Regev *et al*. [1] state that “*To be useful, an atlas must also be an abstraction, comprehensively representing certain features, while ignoring others*.” What then are the appropriate abstractions for a Human Gut Cell Atlas? The answer to this question is guided by what data can be reliably obtained to build the atlas in the first place and then how to map new data onto the atlas, e.g. what is the location from where a resection specimen was obtained, and secondly, what are the questions we want to answer using this data. We start with a simple, clinically orientated, 1D abstraction, which in turn we extend to 2D and 3D models, including the mappings between them. These are complemented with a semantic layer of location descriptions. How we created these abstractions is discussed next and specific details and parameters are provided in the Results section.

### 1D – Core Model

The primary abstraction of the *gut*, representing both the *small* and *large intestines*, is that of a tube connecting the *stomach* to the *anus*. Location is captured in terms of distance along the centreline of the tube to anatomical landmarks, as measured, for example, by the use of an endoscope during a colonoscopy.

### 2D – Anatomograms

Anatomograms have been developed at the European Bioinformatics Institute (EBI) within the Single Cell Expression Atlas (SCEA) programme as 2D graphical representations of certain organs, tissues and cellular assemblies for the purpose of presenting a pictorial overview of transcriptome data and potentially as a graphical interface for data query [17]. Here we have taken the gut 2D anatomogram image and created image domains (regions) that were drawn within the anatomogram for the anatomy of the large and small intestines. These domains were then segmented into sub-domains corresponding to the regions delineated in the anatomogram, e.g. anus, anal canal, descending colon, splenic flexure, etc. Where the anatomogram depicted distant parts of the gut domains as overlapping or touching, e.g. as the small intestine passed behind the transverse colon, then *cut-domains* were created to preserve the appropriate connectivity for the intestines.

Midline-paths were computed from the anus to the tip of the appendix for the large intestine and from the ileocaecal valve – ileum (ICVi) to the gastro-duodenum junction for the small intestine. A propagation algorithm in which possible image locations are considered in priority order was used to compute initial midline paths. Image location priority was determined by a combination of the distance to the path endpoint and from the domain boundary. From the ordered set of path image locations found along each path a smooth B-spline curve was then computed as the primary path representation. All image processing was done using Woolz [18].

### 3D – Radiological-Image Based Models

3D models of the human gut (limited to the large intestine and ileum of the small intestine) were computed from anonymised CT images. Two models have been built, one from an image in which the colon had been inflated and a second from an image in which the colon was non-inflated. In both cases the image domain of the large intestine and all or part of the ileum was segmented from the 3D CT image. For the inflated colon the domain was computed by using threshold-based region growing and morphological operations, with the region growing seed locations entered manually. ITKsnap [19] was used for region growing and Woolz was used for all other image processing operations.

For the non-inflated colon, threshold-based segmentation could not be used because of the wide variation in image values and textures throughout the colonic region, so a pre-segmentation image classification was performed using a machine learning approach based on the full convolutional neural network described by Long *et al* [20] and implemented as “U-Net” by Ronneberger *et al* [21]. To train the convolutional neural network a small number of virtual sections were cut through the 3D image with a range of sectioning parameters – including position and 3D orientation, these were manually segmented using *MAPaint* [22] an interactive drawing application for segmenting 3D image data. The segmented section images (2D) were then used to train a u-net classifier.

To reduce the manual segmentation effort, the number of segmented images was augmented using a combination of affine and non-affine transforms. The trained network was then used to generate a colon classification 2D image for all planes parallel to a virtual section of the original 3D image resulting in a full 3D classification image. The prediction was repeated for 36 sets of virtual sectioning parameters and the resulting 3D classification images were averaged to give a single such image. The classification image was then segmented using region growing and morphological operations in a similar manner to that used for the inflated model. The u-net was built using PyTorch [23], all other image processing was executed using Woolz. With the large intestine and ileum domains segmented from the 3D images, paths through them were computed using the same approach as for the 2D anatomogram.

### Model-Model Mapping Transforms

Each of the 1D, 2D and 3D models represent a spatial context in which data locations can be visualised and queried. It is critical for spatial query and analysis that a location in one model can be mapped to any other so that the spatial frameworks are interoperable and data can be cross compared. For this the 1D linear model was mapped onto the 2D and 3D midline paths computed through the anatomogram and 3D image models respectively. Actual distances along each path are model dependent so a piecewise-linear mapping approach was adopted as an initial or *base-level* cross-mapping. On each path within each of the 1D, 2D and 3D models the landmarks defined in Figure 3 are marked. These are indicated within the anatomogram (Figure 4) and the 3D models (Figure 5) with marker “flags” and a change in colour of the visualised intestine segment. A *location* within a model is defined by the proportional distance along the midline path between the closest proximal (towards the mouth) and closest distal (towards the anus) landmarks. This simple definition allows locations and any data associated with them to be mapped between 1D, 2D and 3D models of the large and small intestines. This base-level mapping of locations between two landmarks without additional information is linear, however, this can be enhanced to a non-linear mapping to better reflect the anatomical structure as more detailed knowledge is acquired. Locations away from the midline path (but within the gut region) are mapped to the closest midline point of the same region. For efficiency this may be precomputed.

**Figure 3:**
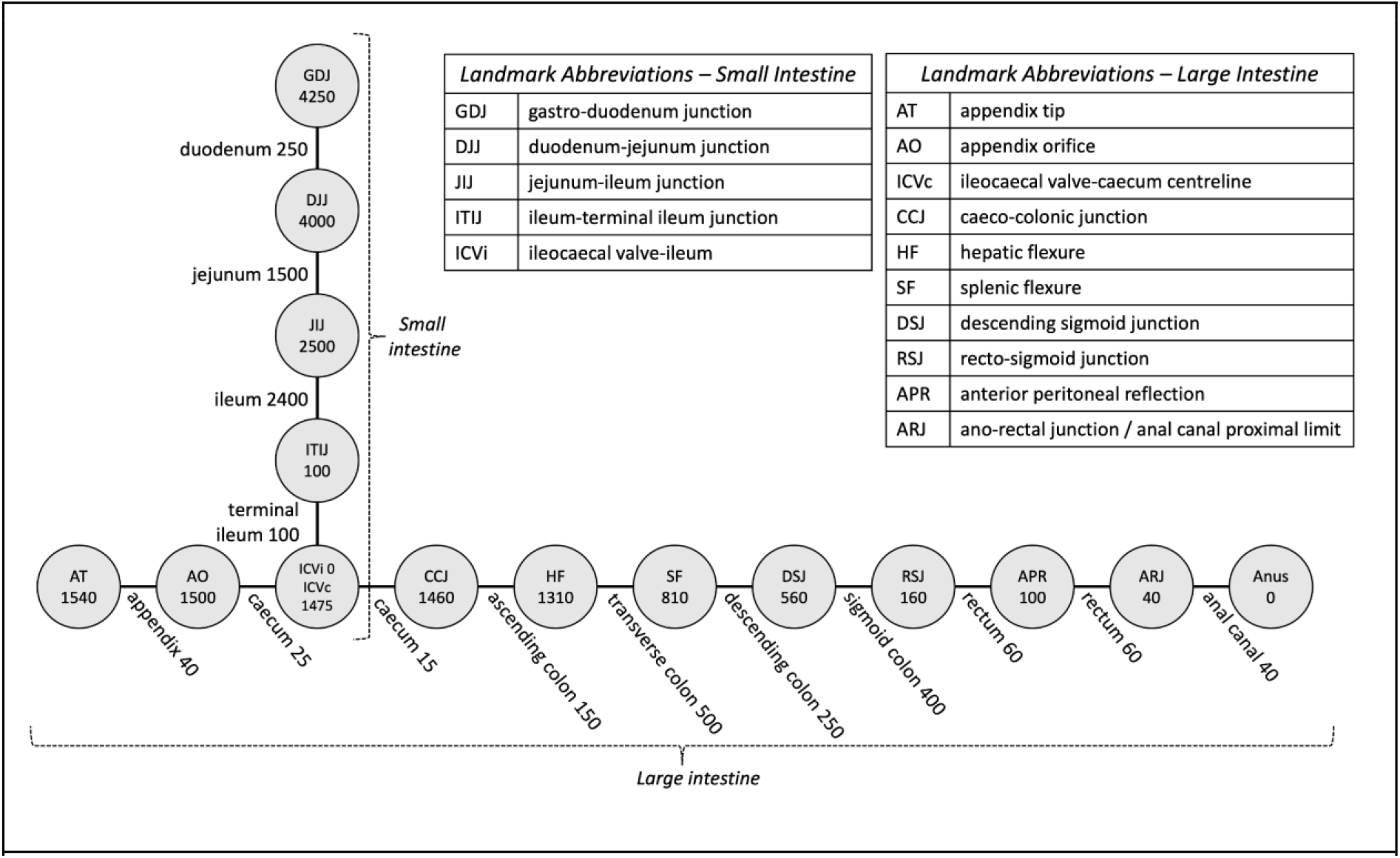
1D Core Model depicted as a graph, with nodes representing anatomical landmarks and edges representing anatomical regions. Regions are generally delimited by their start and end landmarks. The rectum has an additional intermediate landmark known as the anterior peritoneal reflection (APR) and the caecum has an additional intermediate landmark known as the ileocaecal valve (ICV). In the context of the large intestine, the latter is referred to as the ileocaecal valve-caecum centreline (ICVc), whereas the end point for the small intestine is labelled as the ileocaecal valveileum (ICVi). Numbers in nodes represent the landmarks’ distances (in mm) to the anus (for large intestine) and the ICVi (for small intestine). Numbers on links represent the lengths (in mm) of the corresponding gut region. The numbers shown in this diagram reflect a typical live human’s anatomy, different patients will have different measurements.

**Figure 4:**
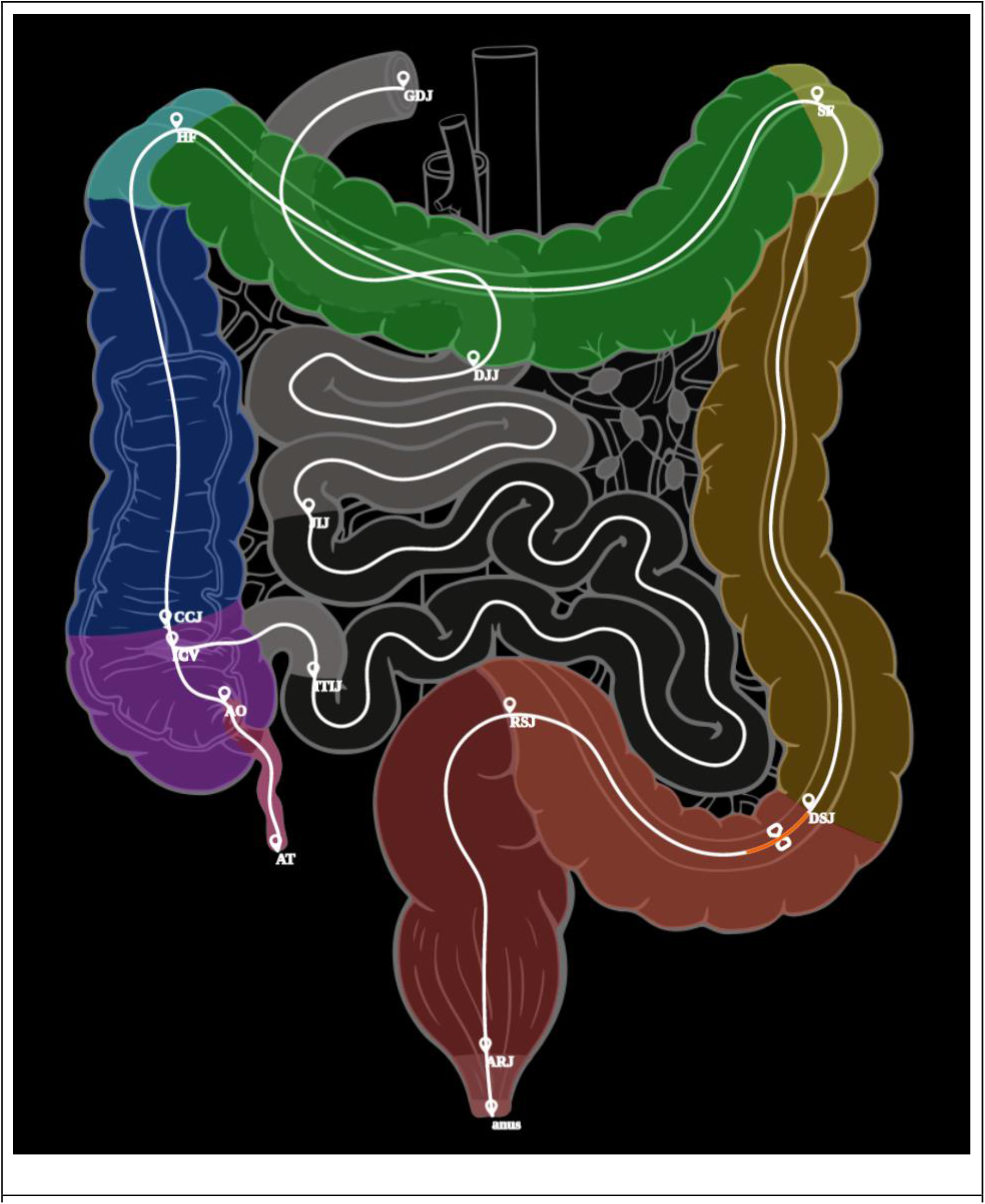
Gut Anatomogram with segmented domains, midline paths and all gut landmarks displayed. This image is captured as a screen shot from the model online visualisation, on the screen the markers and associated text are easier to read. The orange bar at the proximal end of the sigmoid colon depicts the position of the region of interest.

**Figure 5:**
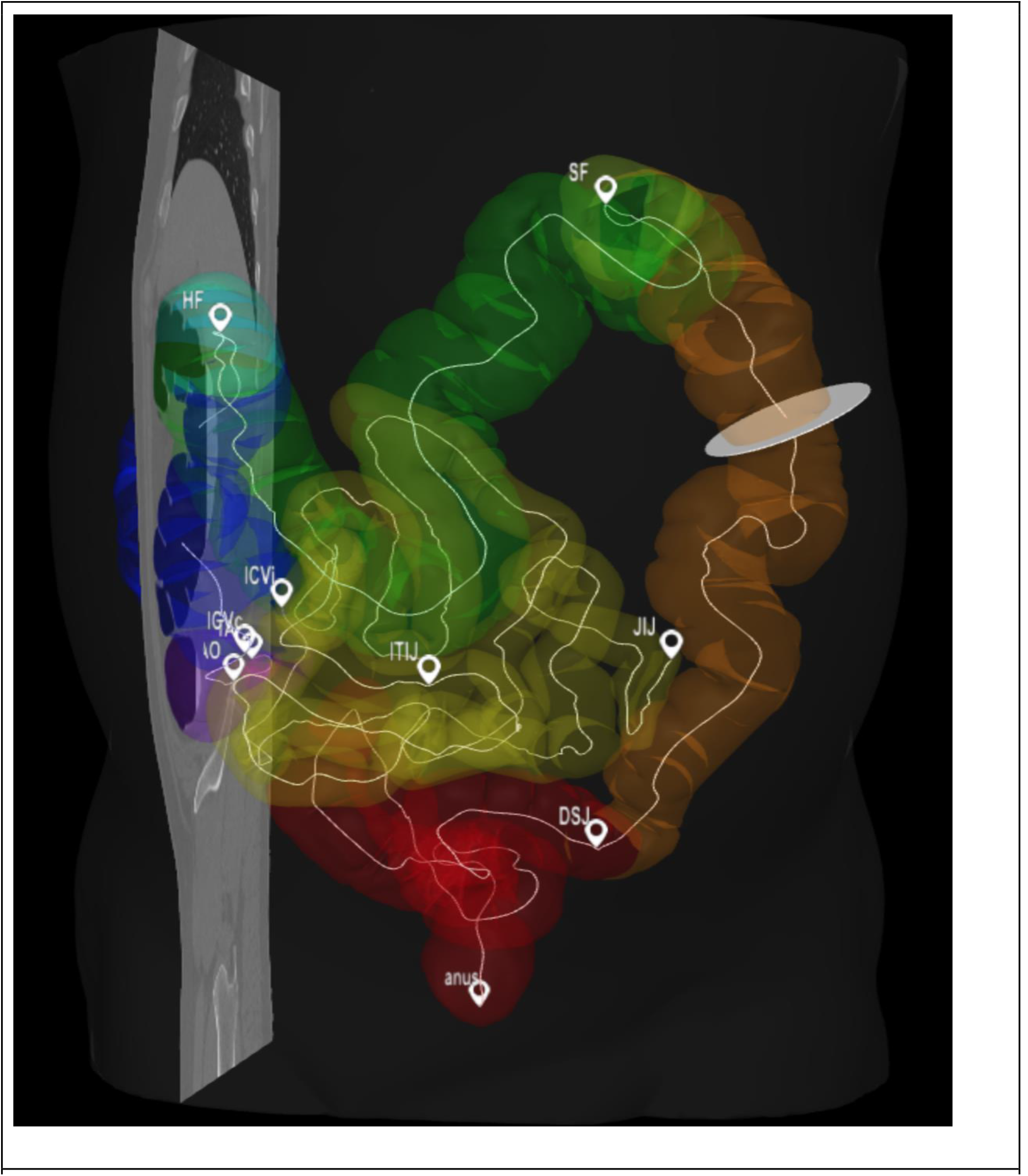
Screenshot of the 3D model viewer showing the segmented large intestine and ileum. The mid-line paths are shown as thin white curved lines with landmarks between sections indicated as small “flags” each labelled with their respective abbreviation. Note in this screenshot image the markers and text are not very distinct compared with the live viewer with a zoom capability. The viewer allows arbitrary sections through the original image to be viewed in the context of the surface models and the cross-sectional disc through the descending colon indicates the position of the image view through the colon shown in Figure 5.

This mapping mechanism allows data from other coordinate frameworks to be mapped to these models. It also allows data from a specific patient to be mapped either through distances to landmarks noted during sample collection or retrospectively using pre-surgery 3D image data.

### Semantic Extension

The descriptions of locations in the gut so far have focused on distances along its tubular structure from the *anus* to the *caecum* and *appendix* (for the *large intestine*) and from the *ileocaecal valve* proximally (for the *small intestine*), but they are unable to provide a mechanism to capture more detailed cell location information with respect to the layers of the gastrointestinal wall at the given position, i.e. the *mucosa, submucosa, muscularis propria* and *serosa*. We therefore semantically complement the location by combining them with the relevant ontological concepts representing these layers, tissue types and cell-types. More specifically, the corresponding standard anatomy terms are those that were agreed as part of the HuBMAP project as a series of Anatomy Structures, Cell Types and Biomarkers (ASCT+B) tables [24, 25], which include links to matching terms in UBERON [26], the Foundational Model of Anatomy (FMA) [27] and the Cell Ontology (CO) [28]. Hence, a typical description might specify the origin of a tissue sample as coming from the *mucosa halfway along the transverse colon* and scRNAseq data could include further specification of cell-type e.g. goblet cell. Future work will also address the representation of location in terms of *villus* vs *crypt* (for *small intestine*) and *left* vs *right* (for *large intestine*).

## Results

### 1D – Core Model

The 1D core model of the gut as described above results in its representation as a graph distinguishing anatomical *landmarks* (nodes) and *regions* (edges), shown in Figure 3. The path from the *anus* to the *caecum* captures the *large intestine*, the path from *terminal ileum* to the *duodenum* captures the *small intestine*. The *ileocaecal valve* node represents the joining point between the *large* and *small intestines*. The numbers attached to nodes represent their respective consensus distance (in mm) to the *anus*, in case of the *large intestine*, and the distance to the *ileocaecal valve* in case of the *small intestine*.

The distances presented here are literature-consensus-based averages [29–31] for an adult living human (distances measured in cadavers are typically longer than the live size due to lack of muscular tone), but of course in reality they vary from person to person. The specific values are provided in Figure 3 and in the model configuration files are available from the project website and associated GitHub archive [32]. Where the actual distances (approximate) for a specific patient can be determined, for example from either a colonoscopic measurement or a CT scan of the patient, a unique model can be derived. The issue of mapping locations across models is discussed below. In the following sections we expand the basic 1D model into 2D and 3D.

### 2D – Anatomograms

The integration of the original gut anatomogram, the segmented domains and the midline paths that we computed for mapping to the other models are shown in Figure 4 and available as a series of configuration and image domain files from the project website and associated GiHub archive [32].

### 3D – Radiological-Image Based Models

The *domains* or 3D regions of the *large intestine* and the *ileum* have been segmented from patient CT images as described above. These have been used to define the boundary surfaces and midline paths through both the *large intestine* from *anus* to *appendix tip* and the *ileum*, from the *ileo-caecal valve* proximally, including *terminal ileum*, for the *small intestine*. Landmarks on these paths have been defined manually using anatomical features visible in the CT scan. Because these features have all been defined with respect to image data then it is possible to display the image data together with this derived data. Figure 5 illustrates one of our patient models showing segmented surfaces, paths from the *anus* to the tip of the *appendix* and from the *ileocaecal valve* to the *ileum* – *jejunum junction*. These are shown with respect to a single virtual plane through the image and a circular virtual section orthogonal to the path through the *descending colon*. Our web-based visualisation allows this derived data including orthogonal sections along the paths to be viewed interactively using a standard web-browser.

The two patient 3D models included so far primarily serve illustrative purposes to help the user of the CCF understand the core concepts of our models in the context of 3D. The longer-term objective is to further automate the methods described earlier to allow the efficient creation of personalised models for each patient, based on their CT and/or MRI scans, that can be mapped into the general Human Gut Cell Atlas CCF.

### Primary Database and Model Json Files and Image Files

The schema concerning the anatomy terms in the Edinburgh GCA database is primarily based on the ASCT+B tables for large and small intestines [25], whose construction has been coordinated by HuBMAP [33] (see also related work below). The data from the ASCT+B resource is parsed and augmented with location information with further additions of specific landmarks which have been identified by the Edinburgh group and which are used in the measurement and mapping of various models. Each model (1D, 2D and 3D) has its own unique configuration file, which is in Json format. The contents of each Json configuration file are generated from the database, which has been further extended to include tables representing each of the Json objects/arrays. The derived configuration file is available from a RESTful web service driven by an Nginx/Payara/ PostgreSQL software stack.

All software and data files are publicly available and available from the Edinburgh Helmsley Gut Cell Atlas web-pages [32] under “Resources”.

### Model Demonstrator Web Application

The Edinburgh Gut Cell Atlas viewer is a web-based application to view and browse a combination of human gut models in 1D, 2D, and 3D. The viewer provides a tool to display locations and map data in the context of the abstract 1D model, the 2D anatomogram model, and 3D gut models reconstructed from volumetric CT scan data. Details of these models and any future model including the anatomical structures, their locations, and other configuration data unique to each model are obtained via an MVC-based RESTful service layer with database connectivity developed as part of the project.

The viewer display consists of two main panels divided by a horizontal splitter as shown in Figure 6. The top panel displays the abstract 1D model and the bottom panel contains the 2D and 3D models. The 1D model view on the top panel is divided into three sub-panels: a slider panel (top), a zoom panel (bottom left), and an additional info panel (bottom right). The slider panel displays the abstract gut model including the colon and the ileum in a linear format with a proportionate scale. The bottom panel of the viewer displays the 2D and 3D models which are organised in a tiled layout by default.

**Figure 6:**
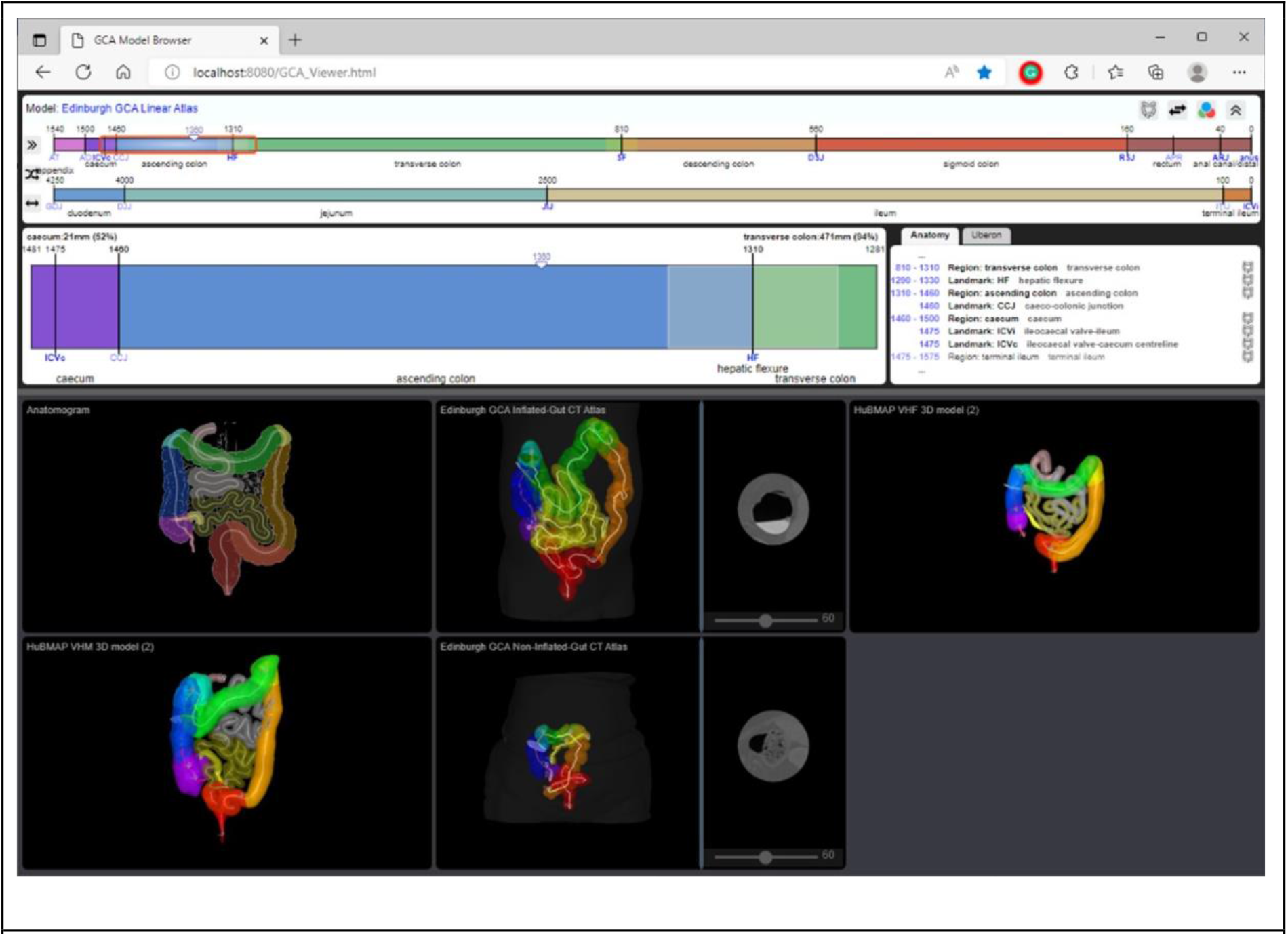
Browser screenshot of Edinburgh Gut Cell Atlas viewer containing the linear model viewer in the upper panel and the 2D and 3D viewer in the lower panel. The browser can be accessed through the project web page [34].

The viewer allows the user to zoom in and zoom out in any of the 2D or 3D models and interactively rotate the 3D models in different directions. Where CT scan image data is associated with a 3D model, image cross-sections orthogonal to the midline paths may be viewed at all locations along the paths. Those section images will display beside the 3D view of the model and are separated by an adjustable vertical splitter. The user can select a subset of the models to be displayed. The viewer allows focus on a 2D or 3D model by maximising the selected model in the bottom panel.

A core concept and a common feature across the models in this viewer is the region of interest (ROI). The ROI is displayed over the 1D model slider as a red rectangle. It specifies the part of the model that is being displayed in the 1D zoom panel and the annotation panel. The 1D zoom panel can potentially be used for adding annotations to the abstract model. The annotation panel provides a textual representation of the 1D model artefacts around the current ROI. Additionally, the panel provides a link to external anatomy data sources including the EBI ontology search (OLS) facility. The ROI is highlighted along the centre line on all of the 2D and 3D models. The extent of the ROI can be changed in the 1D view, and its location can be set on any of the models. Any changes to the location or size of the ROI will be reflected on all the models that are displayed in the viewer.

### Embedding of Interfaces in Applications

The Model Demonstrator software has been designed with reuse in mind. For example, we have prototyped a tool allowing data sets collected in the context of the Helmsley Trust Gut Cell Atlas program to be annotated with Regions of Interest (ROIs). For this a version of the 1D viewer has been embedded in a web user interface. Subsequently, the 1D model viewer can be used to formulate queries according to ROIs (see Figure 7).

**Figure 7:**
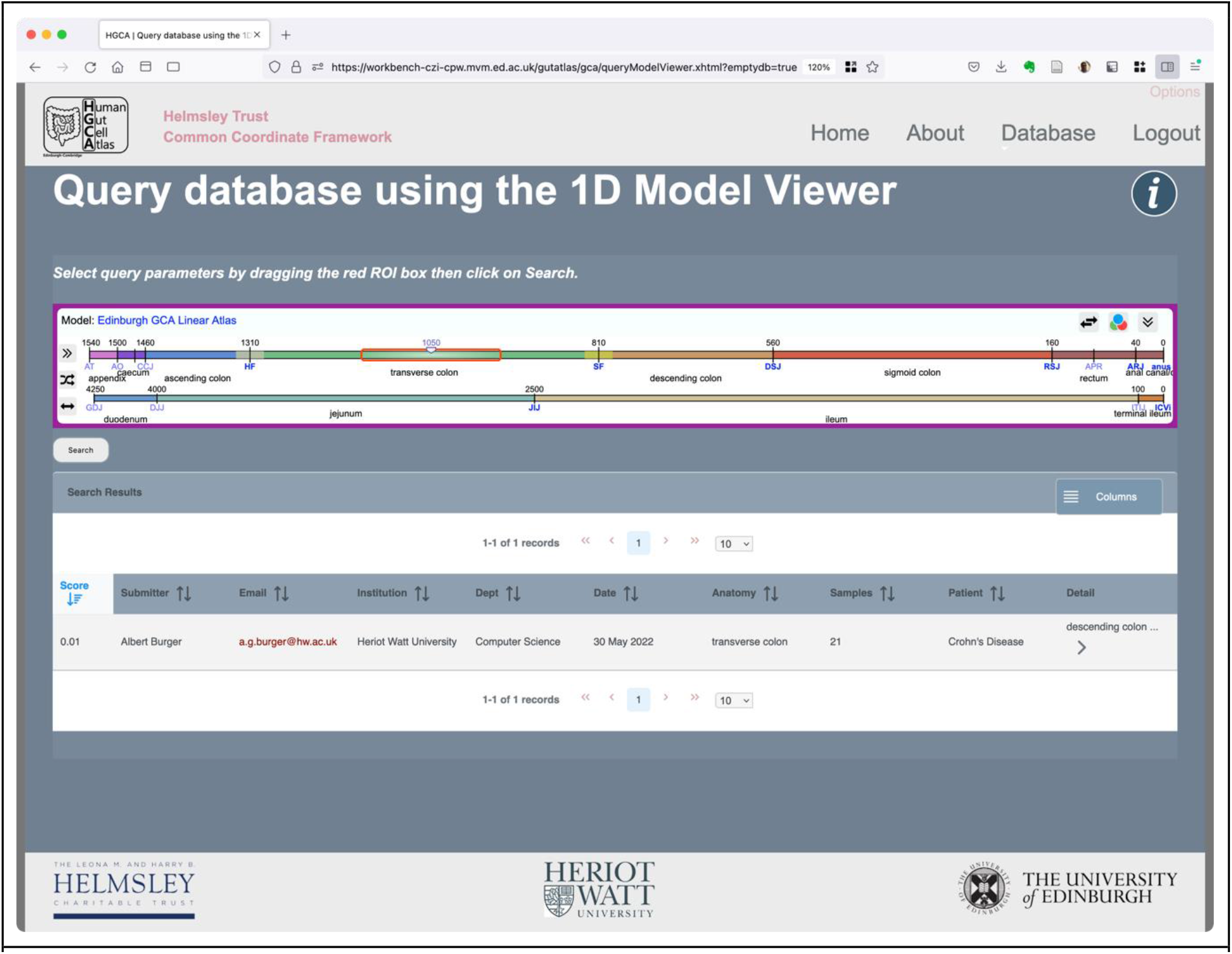
Model-based Query Interface. The red rectangle is used to identify the Region of Interest. Where there is an overlap between the selected query region and the previously recorded image annotation results are returned (sorted by the Jaccard Index).

In addition to the 1D model viewer, the 2D and 3D viewers can similarly be integrated in annotation and query tools.

## Discussion

### Data Resource Integration

This conceptual gut model has been developed to enable spatial annotation of biomedical data from clinical through to single-cell levels of detail. The CCF using this model allows annotation of data in terms of the gut natural coordinate of distance along the gut axis supplemented with anatomy, tissue and cell-type and biomarker ontologies. These allow annotation of data from single-cell analyses through histopathology to macro level clinical data including tissue resections and 3D radiological imaging. Here we present the model with a visualisation tool to allow exploration of the model and to allow the user to define a CCF location as a region-of-interest (ROI) to represent the sample location. We propose that the ROI will represent the spatial location range within which the sample was collected and detailed specification of the sample size will be part of the sample data. Any type of data can include a spatial location defined in this way and we envisage each data resource or archive will make available these locations to allow integration and query in spatial terms. In this way the user can discover relevant data such as single-cell UMAP/t-SNE analyses, histopathology and spatial transcriptomic data linked through to patient MRI data. We are developing a standardised protocol and data format for such spatial annotation for which we coin the term “*location card*”. Any resource using these cards can be cross-queried for data relevant to a particular CCF location with well-defined measures of similarity, proximity and distance to provide data-hits sorted by relevance. In the context of the HCA resource terminology we will be able to implement a spatial query and analysis *data portal*.

The model demonstrator (figure 7) shows how a user can specify a spatial query using the 1D model interface and be presented with an ordered list of entries matching that query. The software provides a simple API to return the detail of the defined ROI and to convert to any of the mapped 2D and 3D models. When the specification for a location card has been defined the API will deliver the card for inclusion as a sample spatial annotation thereby enabling the spatial query capability.

### Current Clinical Practice and Specification of Location

As discussed earlier, endoscope surface distance markings can be used to locate samples (for histology or molecular analyses) within the core linear model described here using distance along the gut centreline. However, there are some limitations that follow from current practice, as some endoscopists only provide the region of the intestines without distance data, and some endoscopists may only provide distance data from the anus, not allowing for individual variation amongst patients in the individual-specific lengths of the intestinal regions. An alternative approach would be to provide sample distance data relative to the nearest landmark (e.g. ileocaecal valve) including the part of the bowel to allow more accurate mapping of sample location within a gut region. This landmark-to-lesion/sample distance data could be provided either as an absolute distance (e.g. ascending colonic lesion located 110mm distal from the ileocaecal valve) or a proportional distance (e.g. ascending colonic lesion located two-thirds of the distance between the ileocaecal valve and the hepatic flexure). This would add to the accuracy of the way the sample location is recorded in clinical records. Use of the 1D centreline and landmark model described here would be able to take advantage of this change, improving the locational accuracy of the data collected and recorded in an appropriate common coordinate framework.

The GCA CCF model viewing system is designed so that samples and numbers of samples can be visualized using all of the 1D, 2D and 3D models in a linked manner. This allows collation of scRNA-Seq, histology and radiology data from both our own research and from other groups in the GCA Consortium, using current demonstrator versions of the model viewer system and we are developing these systems further to allow visualization of sample locations, customised models for specific patients perhaps with resected parts of the intestine or with a stoma. The system permits linking of colonoscopy videos with locations shown in the models. It is important to note that the models are separate from the CCF model visualization software system and either part can be developed or changed independently. The models can be used by other research groups for different research purposes as they wish, using different visualization software.

### Related Frameworks

Although the term *Common Coordinate Framework (CCF)* has only relatively recently (within the last 5 years) become popular, primarily to refer to the computational frameworks required by the emerging Human Cell Atlas (HCA), many of the features of CCFs have been studied and developed in the area of biomedical atlases for much longer. For example, our own work on the Edinburgh Mouse Atlas Project [35] and its coordinate framework was described as early as 1992 [36]. Other biomedical atlases for model organisms include Drosophila [37], chick [38] and zebrafish [39]. Human atlas examples include the HDBR resources [40] and the Allen Human Brain Atlas [41]. A detailed discussion of these atlases is beyond the scope of this paper, but it is noteworthy that just like the newly emerging CCFs, many make use of anatomical ontologies, or at least some form of controlled anatomy vocabulary, and 3D volume reconstructions of the organ or organism under consideration. Importantly, as with the gut common coordinate framework described here, mappings between the ontology concepts and the corresponding voxel sets in the 3D coordinate system are typically provided.

With the introduction of new biotechnology, in particular high-throughput scRNA-seq analysis, extended spatial indexing techniques, now broadly referred to as Common Coordinate Frameworks, have emerged. Rood et al [42] summarise the new requirements and discuss various approaches to CCF development, including a hierarchy of four types of CCFs: macro (whole-organ scale), meso (intra-organ regional scale), micro (histological scale) and fine (cellular scale). Our current GCA models range from macro to micro level. Elmentaite *et al*. [13] describe use of single cell transcriptomics to map the changing gastrointestinal cellular landscape throughout life, from the embryo, through childhood to adult gut, including changes in gut lymphoid tissues in disease.

Boerner et al. [43] introduce a CCF for the National Institutes of Health’s (NIH) Human BioMolecular Atlas Program (HuBMAP). HuBMAP is directly relevant to our own work on a Gut Cell Atlas, since it aims to produce CCF-support for the entire human body, including the small and large intestines. In HuBMAP, expert groups for each organ system are collating and extending the required anatomy, tissue and cellular level ontologies and a series of 3D atlas models are being created to capture the 3D location of tissue samples collected for single-cell analysis in addition to histological imaging. The goal is to deliver a standard “base-level” capability for all parts of the adult female and male human body. For many organ systems including the large and small intestines, this capability has been delivered and some data has already been mapped. The registration tool provides an online 3D interface in which a user can manipulate a small cuboid representing the extent of the sample to a specific location within the 3D model. This is then recorded and can be viewed by all using a similar exploration tool. These interfaces are now available and provide an important base-level mechanism, but have a number of problems for a Gut Cell Atlas (GCA). The current HuBMAP SOP to register a tissue sample [44] provides a mechanism to define the tissue sample as a rectangular block enclosing the sample which is then registered with a 3D location using the registration interface. By this means the block acquires a precise location but with no measure of the spatial uncertainty of that placement. Given the way in which tissue is collected the precise location may not be known and comparison with other data could give rise to inconsistency for example in the interpretation of a gene-expression gradient. A second issue is the usability of the tool which requires practice and is unlikely to be used directly by the expert who collects the tissue - the surgeons, endoscopists and clinical pathologists who have the best knowledge of the positioning and the uncertainties of that sample. A third issue is that the gut atlas system does not seem to have a mechanism to measure functionally relevant proximity from one sample to another, as all coordinates are in a 3D space and the looping of the gut renders 3D proximity much less useful than functional proximity within the “natural coordinate” of the gut specifically distance along the centreline.

In contrast, our own models aim to provide a practical GCA CCF that represents the coordinates within the gut itself along its centreline, relative to nearby landmarks, and provides a mechanism for the clinicians and scientists collecting the data to be able to record a precise location if available, or the uncertainty in location as a range or ROI, in a way that matches the clinical acquisition process. Ultimately, we anticipate that data will be collected and mapped using both approaches, and we are therefore actively collaborating with the HuBMAP team to provide suitable interoperability solutions. At this stage we have used our algorithms to map from our models to the HuBMAP models for small and large intestines for both female and male examples [https://hubmapconsortium.github.io/ccf/pages/ccf-3d-reference-library.html]. Hence, we have established spatial interoperability between all models, a location identified in any of the CCF model spaces can be located in any other. These model mappings are part of the web-based demonstrator application discussed earlier.

We note that using the algorithm described earlier, data can be mapped between the 1D, 2D and 3D models and between these models and others such as the HuBMAP CCF. These simply computed mappings are currently limited to range along a path through the colon or ileum, with no consideration of either radial distance from the midline or angle, although in principle our models and mappings could be extended to include these additional parameters.

Our GCA CCF models were designed specifically for more precise mapping of single cell RNA-sequence (scRNA-Seq) data to location along the central intestinal axis. With accumulation of sufficient scRNA-Seq data, this will allow spatial gradient analysis for investigation of how gut cell gene expression patterns change along the linear axis of the small and large intestines, which may reveal new anatomical or physiological features. This development of data integration involving correlation of scRNA-Seq from multiple sources, is one of the strengths of the HCA approach with use of multi-group consortia. Our approach is complementary to the HuBMAP version which provides only an approximate gut region location, but does not provide a distance-based more accurate sample location, as the HuBMAP mechanism for mapping the position of the sample taken from the gut does not have the capability to precisely locate position in terms of distance from key landmarks or the degree of uncertainty associated with this location. This GCA CCF model enhances and extends the current HCA models for the insights that can be gained from single cell RNA expression data. Furthermore, integration of scRNA-Seq data with associated histology data for any one group is straightforward, whereas integration of data from the same or very similar gut locations from several different research groups requires an accurate location-based approach as set out using this model. Hence, this paper focuses on presenting the new GCA CCF model with a discussion about those novel capabilities that are possible using this model, rather than a demonstration of full use of all of these possible functions.

The GCA is not the only example of a CCF focusing on a single organ. Other organ-specific cell atlas work includes brain [45], lung [46], liver [47], and eye [48].^1^ An interesting exploration of a cross-organ CCF solution is based on the vasculature [49]. Similar to the address of a house in terms of its position on a road, the location of cells is described in terms of the nearest blood vessel. A key advantage here is the vasculature’s natural scaling from arteries to arterioles to capillaries (and similarly for veins). The work is in its early stages and its application as a CCF is yet to be tested.

### Multi Resolution Issues

The HGCA and the encompassing HCA will collect and archive data at a huge range of spatial resolutions: resected material 10-1000 mm, tissue blocks 5-20mm, biopsies 3-5mm, tissue samples for SCA 0.5-3mm, functional tissue units [50] and cellular assemblies 20-100µm, histological sections 5-20µm thickness, single cell data, 10-20µm, and potentially sub-cellular transcript data 0.01-1µm. Image-based data will be at mm resolution (radiology) through to sub-micron resolution microscopy. Current spatial-transcriptomics systems routinely capture the full transcriptome at 20-50µm between sampling locations (dots) and the methodology is improving. The challenge to the CCF is to capture and enable comparison of spatial information across all scales from clinical sample locations through histological section positioning and within image alignment of tissues and multicellular structures. Rood et al. describe a diverse range of approaches for different tissues and discuss a CCF for macro-though to micro-scale data. Here we provide a coordinate-based CCF to capture sample location across the range of 1D to 3D gut representations (also providing interoperability) coupled with semantic refinement for positioning within gut substructures and layers. The 1D conceptual model emphasises the “natural coordinate” of the gut defining the proximal-distal axis and therefore provides a mechanism for linking to natural language descriptions of gut location provided by clinical and pathological reports and also as described in the literature. Semantic descriptions of higher resolution spatial locations such as the relative ordering and sequencing of tissue block and histological sections can also be supported using spatial ontologies such as the BSPO [51], but high-resolution mapping between histological section images is not yet supported.

### Model Organisms and their Mappings

The principle of the 1D model framework for small and large intestines set out here for humans may also be used for linear representation of the intestines of other species, including a range of model organisms such as mouse, rat, pig, sheep, etc. Co-linear mapping of the small intestine from gastro-duodenal junction to ileocaecal valve, and similarly the large intestine from ileocaecal valve to anus, across different species allows approximate translation of samples or lesions between similar regions of mammalian gut. This would allow biomedically meaningful spatial mappings between these different species in order to facilitate cross-species data comparison, including support for search and query functionality across many different species. This may form the basis of a Cross-Species Intestinal Cell Atlas for development of standards and comparisons around intestinal data.

A detailed discussion and evaluation of this cross-species work is beyond the scope of this paper. However, we note the potential utility of the linear model approach in this context. Figure 8 shows the use of an *abstract* model as an intermediate, not unlike UBERON [26] as a cross-species anatomy ontology. Although we could map directly between mouse and human, the abstract model simplifies the addition of other species. It only requires mapping each species onto the abstract model instead of all possible pairwise combinations of species.

**Figure 8:**
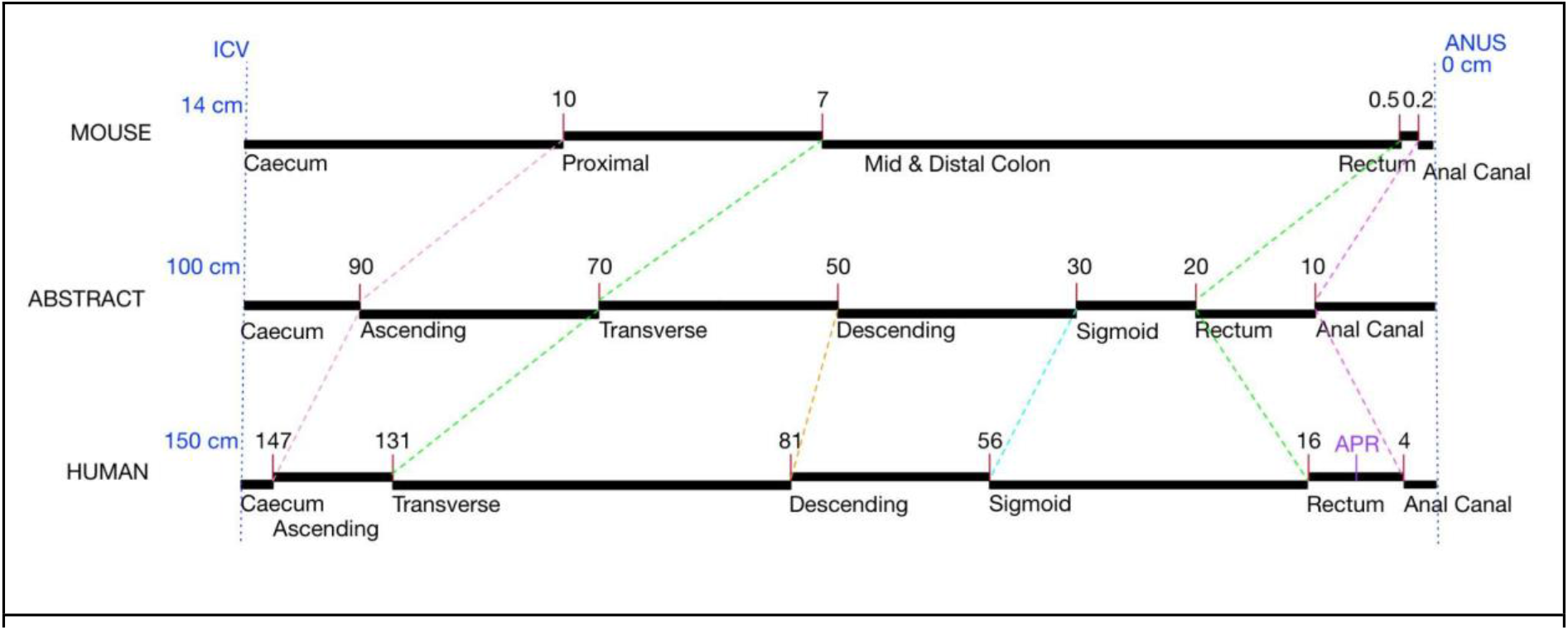
Linear Model-based Mapping of Gut Location across mouse and human using an abstract intermediate.

## Conclusions

Here, we conclude that the small and large intestines have a natural “gut coordinate” system that is best represented as a 1D centreline through the gut tube. This reflects the functional differences that arise in different regions along the small and large intestines, whilst deliberately ignoring the three-dimensional looping of the intestines which may generate transient and spurious proximity between widely spaced and very different regions of gut. We have constructed a 1D model, based on the centreline of small and large intestines, with accompanying viewer software that allows visualisation of this 1D representation, along with interoperable translation to both a 2D anatomogram model and multiple 3D radiological scan-based models of the intestines. This permits users to locate samples (taken for histology, single cell transcriptomics, or other research purposes) or clinically identified lesions from endoscopy or radiological investigations or surgical intervention, to be accurately located using either absolute distance or proportional distance along the centreline from one or more known landmarks. Importantly, this mechanism for more specific anatomical location of samples allows data comparison between different patients and different studies in a functionally relevant manner.

## Abbreviations

ASCT+B: Anatomical Structures, Cell Types, plus Biomarkers
BSPO: Biological Spatial Ontology
CCF: Common Coordinate Framework
CD: Crohn’s Disease
CO: Cell Ontology
CT: Computed Tomography
EBI: European Bioinformatics Institute
FMA: Foundational Model of Anatomy
HCA: Human Cell Atlas
HDBR: Human Developmental Biology Resource HGCA Human Gut Cell Atlas
HuBMAP: Human BioMolecular Atlas Program IBD Inflammatory Bowel Disease
ICV: Ileocaecal Valve
JSON: JavaScript Object Notation
MAST: Model-based Analysis of Single-cell Transcriptomics
MRI: Magnetic Resonance Imaging
MVC: Model-View-Controller
OLS: Ontology Lookup Service
PCA: Principal Component Analysis
RNA: Ribonucleic Acid
ROI: Region Of Interest
SCA: Single Cell Analysis
SCEA: Single Cell Expression Atlas
t-SNE: t-distributed stochastic neighbour embedding
UMAP: Uniform Manifold Approximation and Projection
UMI: Unique Molecular Identifier

## Declarations

### Ethics approval and consent to participate

Image data used for the radiological scan-based 3D models of small and large intestines were fully anonymised datasets that were made available in accordance with the appropriate UK ethical and legal framework.

### Consent for publication

Not applicable.

### Availability of data and materials

All the model configuration data, image data and software is fully open-source and freely available from the Edinburgh HGCA web-site (https://www.ed.ac.uk/comparative-pathology/the-gut-cell-atlas-project/project-resources) as a compressed archive file as well as from a GitHub repository (https://github.com/orgs/Comparative-Pathology/repositories) referenced from that site.

### Authors’ contributions

MA is lead-PI of the project, AB, RB, DA, SD and IP are co-PIs. All authors have contributed to the conceptual model work. BH developed the imaging software. DH, MS and MW developed the web-based user interfaces and backend databases. MG carried out the medical informatics analysis. All authors have contributed to the manuscript. AB coordinated the authoring efforts. All authors have read and approved the manuscript.

## Acknowledgements

The authors wish to acknowledge constructive discussions with Kenneth McLeod, previously at Heriot-Watt University, regarding the 1D framework and model organisms; regular discussions about development of the common coordinate framework with both Jessica Langer and Stephanie LeValley of The Leona M. and Harry B. Helmsley Charitable Trust; and early funding for related work by the Chan-Zuckerberg Initiative during which the 1D linear gut model emerged. Dr Stephen Glancy provided the anonymised radiological images. SD acknowledges the support of NHS Research Scotland via NHS Lothian.

## Footnotes

1 Please note that these are just examples and do not constitute a comprehensive list of all existing cell atlases and CCFs.

## References

1. Regev A, Teichmann SA, Lander ES, Amit I, Benoist C, Birney E, et al. The Human Cell Atlas. eLife. 2017;6:e27041.

2. Human Cell Atlas Home Page. https://www.humancellatlas.org/. Accessed 18 Nov 2022.

3. Alatab S, Sepanlou SG, Ikuta K, Vahedi H, Bisignano C, Safiri S, et al. The global, regional, and national burden of inflammatory bowel disease in 195 countries and territories, 1990–2017: a systematic analysis for the Global Burden of Disease Study 2017. Lancet Gastroenterol Hepatol. 2020;5:17–30.

4. Narula N, Wong ECL, Dehghan M, Mente A, Rangarajan S, Lanas F, et al. Association of ultra-processed food intake with risk of inflammatory bowel disease: prospective cohort study. BMJ. 2021;374:n1554.

5. Racine A, Carbonnel F, Chan SSM, Hart AR, Bueno-de-Mesquita HB, Oldenburg B, et al. Dietary Patterns and Risk of Inflammatory Bowel Disease in Europe: Results from the EPIC Study. Inflamm Bowel Dis. 2016;22:345–54.

6. Xia B, Yang M, Nguyen LH, He Q, Zhen J, Yu Y, et al. Regular Use of Proton Pump Inhibitor and the Risk of Inflammatory Bowel Disease: Pooled Analysis of 3 Prospective Cohorts. Gastroenterology. 2021;161:1842-1852.e10.

7. Jones G-R, Lyons M, Plevris N, Jenkinson PW, Bisset C, Burgess C, et al. IBD prevalence in Lothian, Scotland, derived by capture-recapture methodology. Gut. 2019;68:1953–60.

8. Lamb CA, Kennedy NA, Raine T, Hendy PA, Smith PJ, Limdi JK, et al. British Society of Gastroenterology consensus guidelines on the management of inflammatory bowel disease in adults. Gut. 2019;68 Suppl 3:s1–106.

9. Lennard-Jones JE, Shivananda S. Clinical uniformity of inflammatory bowel disease a presentation and during the first year of disease in the north and south of Europe. EC-IBD Study Group. Eur J Gastroenterol Hepatol. 1997;9:353–9.

10. Gajendran M, Loganathan P, Catinella AP, Hashash JG. A comprehensive review and update on Crohn’s disease. Dis--Mon DM. 2018;64:20–57.

11. Alfredsson J, Wick MJ. Mechanism of fibrosis and stricture formation in Crohn’s disease. Scand J Immunol. 2020;92:e12990.

12. Moreno P, Fexova S, George N, Manning J, Miao Z, Mohammed S, et al. Expression Atlas update: gene and protein expression in multiple species. Nucleic Acids Res. 2021;50.

13. Elmentaite R, Kumasaka N, Roberts K, Fleming A, Dann E, King HW, et al. Cells of the human intestinal tract mapped across space and time. Nature. 2021;597:250–5.

14. Jovic D, Liang X, Zeng H, Lin L, Xu F, Luo Y. Single-cell RNA sequencing technologies and applications: A brief overview. Clin Transl Med. 2022;12:e694.

15. Common Coordinate Framework (CCF) Meeting. 2017.

16. Goldacre B, Morley J. Better, broader, safer: using health data for research and analysis. Department of Health and Social Care; 2022.

17. Moreno P, Fexova S, George N, Manning JR, Miao Z, Mohammed S, et al. Expression Atlas update: gene and protein expression in multiple species. Nucleic Acids Res. 2022;50:D129–40.

18. ma-tech. ma-tech/Woolz. 2021.

19. ITK-SNAP Home. http://www.itksnap.org/pmwiki/pmwiki.php. Accessed 16 Jan 2022.

20. Long J, Shelhamer E, Darrell T. Fully convolutional networks for semantic segmentation. In: 2015 IEEE Conference on Computer Vision and Pattern Recognition (CVPR). 2015. p. 3431–40.

21. Ronneberger O, Fischer P, Brox T. U-Net: Convolutional Networks for Biomedical Image Segmentation. In: Navab N, Hornegger J, Wells WM, Frangi AF, editors. Medical Image Computing and Computer-Assisted Intervention – MICCAI 2015. Cham: Springer International Publishing; 2015. p. 234–41.

22. ma-tech. ma-tech/MAPaint. 2020.

23. PyTorch. https://www.pytorch.org. Accessed 16 Jan 2022.

24. Herr BW, Hardi J, Quardokus EM, Bueckle A, Chen L, Wang F, et al. Specimen, Biological Structure, and Spatial Ontologies in Support of a Human Reference Atlas. 2022;:2022.09.08.507220.

25. Börner K, Teichmann SA, Quardokus EM, Gee J, Browne K, Osumi-Sutherland D, et al. Anatomical Structures, Cell Types, and Biomarkers Tables Plus 3D Reference Organs in Support of a Human Reference Atlas. 2021;:2021.05.31.446440.

26. Haendel MA, Balhoff JP, Bastian FB, Blackburn DC, Blake JA, Bradford Y, et al. Unification of multi-species vertebrate anatomy ontologies for comparative biology in Uberon. J Biomed Semant. 2014;5:21.

27. Rosse C, Mejino JLV. A reference ontology for biomedical informatics: the Foundational Model of Anatomy. J Biomed Inform. 2003;36:478–500.

28. Diehl AD, Meehan TF, Bradford YM, Brush MH, Dahdul WM, Dougall DS, et al. The Cell Ontology 2016: enhanced content, modularization, and ontology interoperability. J Biomed Semant. 2016;7:44.

29. Treuting PM, Arends MJ, Dintzis SM. 11 - Upper Gastrointestinal Tract. In: Treuting PM, Dintzis SM, Montine KS, editors. Comparative Anatomy and Histology (Second Edition). San Diego: Academic Press; 2018. p. 191–211.

30. Treuting PM, Arends MJ, Dintzis SM. 12 - Lower Gastrointestinal Tract. In: Treuting PM, Dintzis SM, Montine KS, editors. Comparative Anatomy and Histology (Second Edition). San Diego: Academic Press; 2018. p. 213–28.

31. Tortora GJ. Principles of anatomy and physiology / Gerard J. Tortora, Sandra Reynolds Grabowski. 10th ed. New York: J. Wiley & Sons; 2003.

32. The Helmsley Gut Cell Atlas Project. The University of Edinburgh. https://www.ed.ac.uk/comparative-pathology/the-gut-cell-atlas-project. Accessed 26 Jun 2022.

33. Snyder MP, Lin S, Posgai A, Atkinson M, Regev A, Rood J, et al. The human body at cellular resolution: the NIH Human Biomolecular Atlas Program. Nature. 2019;574:187–92.

34. Gut Atlas Models. The University of Edinburgh. https://www.ed.ac.uk/comparative-pathology/the-gut-cell-atlas-project/project-resources/gut-atlas-models. Accessed 26 Jun 2022.

35. Armit C, Richardson L, Hill B, Yang Y, Baldock RA. eMouseAtlas informatics: embryo atlas and gene expression database. Mamm Genome. 2015;26:431–40.

36. Baldock R, Bard J, Kaufman M, Davidson D. A real mouse for your computer. BioEssays News Rev Mol Cell Dev Biol. 1992;14:501–2.

37. Gramates LS, Agapite J, Attrill H, Calvi BR, Crosby MA, dos Santos G, et al. FlyBase: a guided tour of highlighted features. Genetics. 2022;220:iyac035.

38. Wong F, Welten MCM, Anderson C, Bain AA, Liu J, Wicks MN, et al. eChickAtlas: An introduction to the database. genesis. 2013;51:365–71.

39. Bradford YM, Van Slyke CE, Ruzicka L, Singer A, Eagle A, Fashena D, et al. Zebrafish information network, the knowledgebase for Danio rerio research. Genetics. 2022;220:iyac016.

40. Kerwin J, Yang Y, Merchan P, Sarma S, Thompson J, Wang X, et al. The HUDSEN Atlas: a three-dimensional (3D) spatial framework for studying gene expression in the developing human brain. J Anat. 2010;217:289–99.

41. Hawrylycz MJ, Lein ES, Guillozet-Bongaarts AL, Shen EH, Ng L, Miller JA, et al. An anatomically comprehensive atlas of the adult human brain transcriptome. Nature. 2012;489:391–9.

42. Rood JE, Stuart T, Ghazanfar S, Biancalani T, Fisher E, Butler A, et al. Toward a Common Coordinate Framework for the Human Body. Cell. 2019;179:1455–67.

43. Börner K, Quardokus EM, Herr II BW, Cross LE, Record EG, Ju Y, et al. Construction and Usage of a Human Body Common Coordinate Framework Comprising Clinical, Semantic, and Spatial Ontologies. ArXiv200714474 Cs Q-Bio. 2020.

44. Bueckle A. SOP: Using the CCF Registration User Interface. 2022. https://doi.org/10.5281/zenodo.6628366.

45. Eze UC, Bhaduri A, Haeussler M, Nowakowski TJ, Kriegstein AR. Single-cell atlas of early human brain development highlights heterogeneity of human neuroepithelial cells and early radial glia. Nat Neurosci. 2021;24:584–94.

46. Ardini-Poleske ME, Clark RF, Ansong C, Carson JP, Corley RA, Deutsch GH, et al. LungMAP: The Molecular Atlas of Lung Development Program. Am J Physiol - Lung Cell Mol Physiol. 2017;:ajplung.00139.2017.

47. Aizarani N, Saviano A, Sagar Mailly L, Durand S, Pessaux P, et al. A Human Liver Cell Atlas: Revealing Cell Type Heterogeneity and Adult Liver Progenitors by Single-Cell RNA-sequencing. bioRxiv. 2019. https://doi.org/10.1101/649194.

48. Gautam P, Hamashima K, Chen Y, Zeng Y, Makovoz B, Parikh BH, et al. Multi-species single-cell transcriptomic analysis of ocular compartment regulons. Nat Commun. 2021;12:5675.

49. Weber GM, Ju Y, Börner K. Considerations for Using the Vasculature as a Coordinate System to Map All the Cells in the Human Body. Front Cardiovasc Med. 2020;7.

50. de Bono B, Grenon P, Baldock R, Hunter P. Functional tissue units and their primary tissue motifs in multi-scale physiology. J Biomed Semant. 2013;4:22.

51. Dahdul WM, Cui H, Mabee PM, Mungall CJ, Osumi-Sutherland D, Walls RL, et al. Nose to tail, roots to shoots: spatial descriptors for phenotypic diversity in the Biological Spatial Ontology. J Biomed Semant. 2014;5:34.

